# Protein Kinases MpkA and SepH Transduce Crosstalk Between CWI and SIN Pathways to Activate Protective Hyphal Septation Under Echinocandin Cell Wall Stress

**DOI:** 10.1101/2024.07.23.604799

**Authors:** Alexander G. Doan, Jessica E. Schafer, Casey M. Douglas, Matthew S. Quintanilla, Meredith E. Morse, Harley Edwards, Walker D. Huso, Kelsey J. Gray, JungHun Lee, Joshua K. Dayie, Steven D. Harris, Mark R. Marten

**Affiliations:** Department of Chemical, Biochemical, and Environmental Engineering, University of Maryland, Baltimore County; Department of Plant Pathology, Entomology, and Microbiology, Iowa State University

## Abstract

This study investigates a previously unreported stress signal transduced as crosstalk between the Cell Wall Integrity (CWI) pathway and the Septation Initiation Network (SIN). Echinocandins, which target cell wall synthesis, are widely used to treat mycoses. Their efficacy, however, is species specific; our findings suggest this is due largely to CWI-SIN crosstalk and the ability of filamentous species to fortify with septa in response to echinocandin stress. To better understand this crosstalk, we used a microscopy-based assay to measure septum density, aiming to understand the septation response to cell wall stress. The echinocandin micafungin, an inhibitor of β-(1,3)-glucan synthase, was employed to induce this stress. We observed a strong positive correlation between micafungin treatment and septum density in *wild-type* strains. This finding suggests that CWI activates SIN under cell wall stress, increasing septum density to protect against cell wall failure. More detailed investigations, with targeted knockouts of CWI and SIN signaling proteins, enabled us to identify crosstalk occurring between the CWI kinase, MpkA, and the SIN kinase, SepH. This discovery, of previously unknown crosstalk between the CWI and SIN pathways, not only reshapes our understanding of fungal stress responses, but also unveils a promising new target pathway for the development of novel antifungal strategies.

**IMPORTANCE:** Echinocandin resistant species pose a major challenge in clinical mycology by rendering one of only four lines of treatment, notably one of the two that are well-tolerated, ineffective in treating systemic mycoses of these species. Previous studies have demonstrated that echinocandins fail against highly polarized fungi because they target only apical septal compartments. It is known that many filamentous species respond to cell wall stress with hyperseptation. In this work we show that echinocandin resistance hinges on this dynamic response, rather than on innate septation alone. We also unveil, for the first time, the signaling pathway used to deploy the hyperseptation response. By disabling this pathway, we were able to render mycelia susceptible to echinocandin stress. Hence, this work enhances our microbiological understanding of filamentous fungi and introduces a potential target to overcome echinocandin resistant species.

## BACKGROUND

Fungi comprise a diverse group of species capable of living in a wide range of environments, including as pathogens in both animal and plant tissues^1,2^. Notable human pathogens include *Aspergillus fumigatus*, *Candida albicans*, *Cryptococcus spp.*, and *Mucor spp.*^3^, which are responsible for a majority of the 1.5 million deaths caused by fungal infections annually^4^. Understanding the molecular mechanisms that underlie these pathogens’ ability to respond to stress is crucial for elucidating their virulence and developing strategies to mitigate their pathogenicity.

To date, there are only four lines of treatment available for systemic mycoses: azoles, echinocandins, amphotericin B, and flucytosine in combination with amphotericin B, all of which target either the fungal cell wall or membrane and have species-specific efficacy ^5^. The high mortality of systemic mycoses^4^ combined with the advancement of drug-resistant strains presents a dire need to develop new lines of treatment^5^.

Although many of the mechanisms we discuss are conserved broadly, for brevity we use *Aspergillus* names for genes and proteins in this paper and provide the names of orthologous proteins in Table 1 for the reader to compare with other common ascomycete models.

**Table 1.** Orthology of signaling proteins described in this work for common model ascomycetes. BH = Blast Best Hit.

| *A. nidulans* | *S. pombe* | *S. cerevisiae* | *N. crassa* |
| --- | --- | --- | --- |
| AnkA | Wee1 | SWE1 | STK-19 |
| BckA | Mkh1 (BH) | BCK1 | MIK-1 |
| BubA | Cdc16 | BUB2 | DIV-2 |
| ByrA | Byr4 |  | NCU01584 |
| MkkA | Pek1 | MKK1, MKK2 | MEK-1, MEK-2 |
| MobA | Mob1 | MOB1 | MOB-1 |
| MpkA | Pmk1, Spk1 | SLT2, KSS1, FUS3 | MAK-1, MAK-2 |
| NimX | Cdc1/Cdc2, Pef1 | CDC28, PHO85 | CDC28, MDK-1 |
| PkcA | Pck1, Pck2 | PKC1 | PKC |
| PomA | Pom1, Pom2, Ppk15 | YAK1 | PRK-2, STK-46 |
| RlmA | Mbx2 | SMP1 | TCF-23 |
| SepH | Cdc7 | CDC15 | CDC15 |
| SepL | Ppk11, Sid1 | SPS1, KIC1 | PRK-9, STK-6 |
| SepM | Cdc14 |  | DIV-36 |
| SidB | Sid2 | DBF2, DBF20 | DBF-2 |
| SpgA | Spg1 | TEM1 | DIV-17 |

The fungal cell wall is critical to viability and pathogenicity^6^ and is targeted by echinocandins through inhibition of β-(1,3)-glucan synthase^5^, which is essential for the formation of the major cell wall structural component β-(1,3)-glucan^6,7^. Fungi are able to respond to echinocandin-induced wall stress through the Cell Wall Integrity (CWI or CWIS) pathway. CWI works to maintain integrity of the cell wall in response to cell wall stress and is highly conserved in the fungal kingdom^7–10^. The CWI pathway is activated when wall stress triggers cell surface receptors^11^. These receptors activate the guanine nucleotide exchange factor (GEF), Rom2, which activates the guanine nucleotide binding protein (G-protein) RhoA^9,12^. RhoA then phosphorylates PkcA kinase^11,13–16^, which initiates the CWI mitogen activated protein kinase (MAPK) cascade BckA-MkkA-MpkA^17–20^. MpkA is the terminal kinase of CWI and activates the transcription factor RlmA to regulate cell wall biosynthesis genes transcriptionally^9,19,21^. MpkA also activates other proteins directly through protein-protein interactions^18,22^.

Filamentous fungi are able to survive echinocandin-induced wall stress through the construction of septa^23–25^, which are structures used to partition hyphae into multiple compartments^26–34^. The compartmentalization of hyphal cytoplasm by septa protects germlings from a complete loss of cytoplasm in the event that the cell wall fails to contain the germling’s turgor pressure^29,35–38^. Nominally, septa allow for free flow of cytoplasm between compartments^39,40^. However, if hyphae suffer a cell wall rupturing injury, Woronin bodies migrate to the septal pore and serve to block cytoplasm flow from healthy hyphal compartments into the injured one, thereby saving the healthy septal compartments^31,34,41,42^.

Echinocandins, such as micafungin (MF), target active β-(1,3)-glucan synthesis^43,44^. For species that grow with a high degree of polarity, e.g. *Aspergillus spp*., the echinocandin drug effect occurs primarily at the hyphal tip, where there is active β-(1,3)-glucan synthesis^5,45^. Non-apical septal compartments are protected from echinocandin wall stress by the separation septa provide between them and the more vulnerable apical compartments. Hence, echinocandins result in a fungistatic effect for *Aspergillus spp.*^5,43,45^, as subapical compartments are largely unaffected^46^.

Septation in filamentous fungi is regulated by the Septation Initiation Network (SIN)^26,30,47,48^ (MEN in yeast^49^ and HIPPO in higher eukaryotes^50^). Although the sources of all septation signals have not been mapped, one major component of SIN is the protein kinase SepH^24–26,28,30,47,51–55^. When activated, SepH initiates septation through the kinase cascade SepH-SepL-SidB, where SepL and SidB are dependent on cofactors SepM and MobA, respectively^24,25,28,47,52,53^. SepH has several known upstream regulators. The cyclin dependent protein kinase NimX regulates SepH through the small GTPase SpgA and its GTPase-activating proteins (GAPs) ByrA and BubA^24–26,30,47,56^. Negative regulators of SIN have also been reported. The protein kinase PomA has roles regulating cell polarity in *S. pombe*^57–60^ and has been shown to be a negative regulator of SIN activity in *A. nidulans*^52^. The cell cycle regulator, AnkA^59,61–71^, another protein kinase, has also been shown to repress SIN activity^52^.

Mycelia lacking genes required to form septa have been shown to survive at significantly lower rates compared to their *wild-type* counterparts when exposed to critical concentrations of cell wall perturbing drugs^23–25^. Control experiments carried out by Spence et al. (2022) also determined that survival rates of non-septating strains were not affected by critical concentrations of fungicidal drugs that do not target the cell wall^23^. These results imply that septa are critical specifically to survive cell wall stress.

These findings, in combination with evidence from our previous work (showing that SIN is activated by the CWI pathway in response to micafungin-induced cell wall stress)^18^, have led us to investigate this interaction. The CWI-SIN interaction could represent a critical adaptive mechanism that allows vegetive hyphae to rapidly compartmentalize cytoplasm in order to minimize losses under increased risk of cell wall failure. The potential of such a mechanism opens avenues for novel therapeutic strategies that could disable this protective feature when targeting the fungal cell wall, e.g. with echinocandins.

This work aims to elucidate the molecular dialogue between the CWI and SIN signaling pathways through genetic means. As *Aspergillus nidulans* has proven to be a productive model organism, due to its genetic tractability^72^, its close relation to the pathogen *Aspergillus fumigatus*^73^, and the fact that it features many conserved pathways common amongst many other fungal species^74–78^, we have chosen to conduct our studies in this model.

## RESULTS

### SIN is Activated by Micafungin Cell Wall Stress

In our previous study, Chelius et al. (2020)^18^, we found that under micafungin-induced cell wall stress, there was an increase in septum density (septa/µm^2^ growth area), and the SIN kinase SidB was differentially phosphorylated at three phosphorylation sites. We reproduced the septum density experiments with the *wild-type* control strain (FGSC^79^ A1405) in more detail, and the results have been summarized in Figure 1.

**Figure 1.**
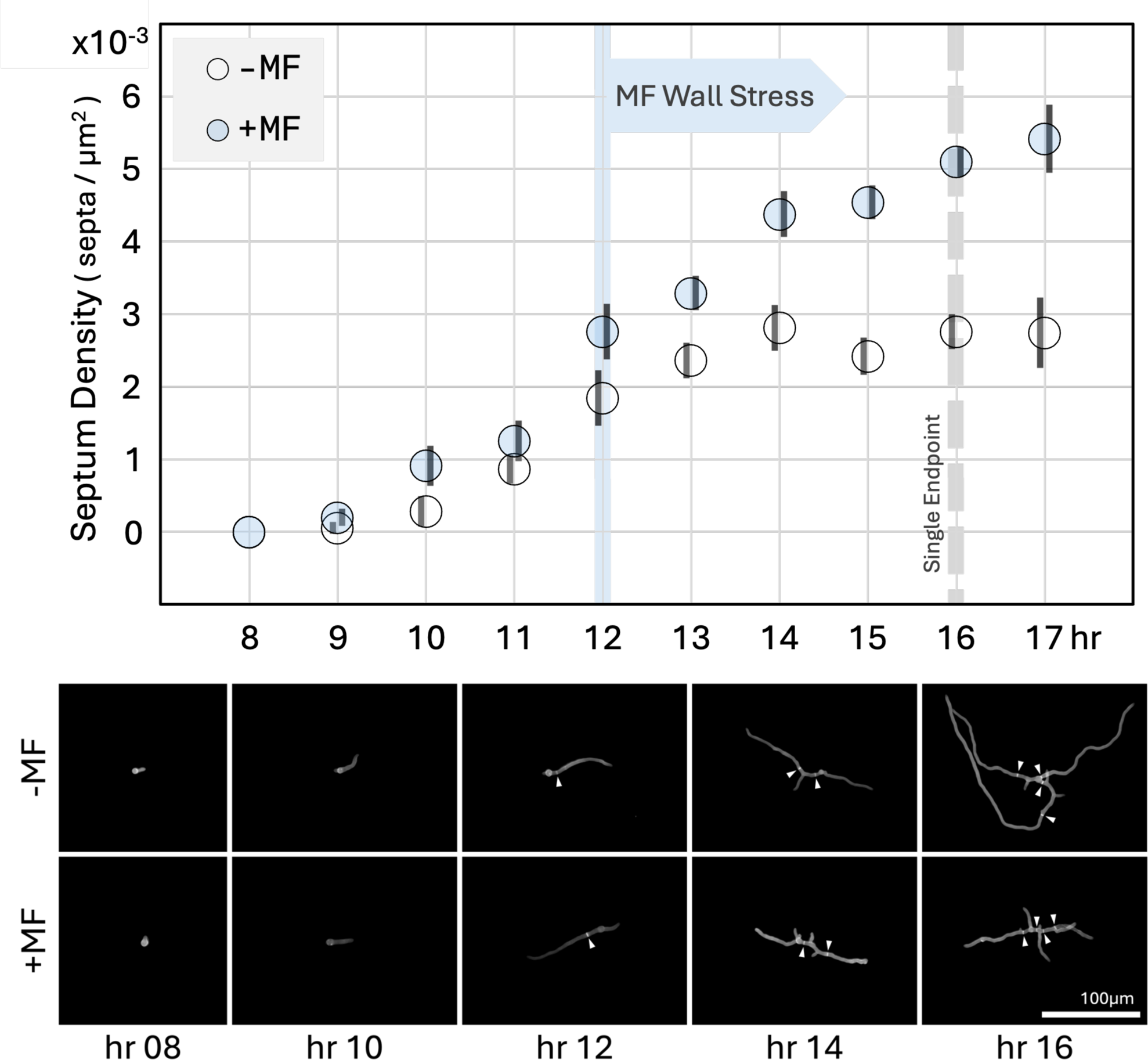
<u>Top</u>: Time-course of septum density from 8-17 hours after coverslip inoculation. Freshly harvested spores were adhered to coverslips using concanavalin A and then submerged in liquid YGV growth media at 28°C. Micafungin (MF) was added for the shaded series 12 hours after inoculation (blue flag) to a final concentration of 10 ng/mL. For each marker n ≈ 90. Single endpoint data for Figure 2 was collected at dashed line. Error bars are standard error. <u>Bottom</u>: Selected images of germlings represented by plot data. White arrowheads highlight septa.

For the shaded series (+MF), we observed a dramatic increase in septum density starting when germlings were exposed to 10 ng/mL micafungin starting 12 hours after inoculation, which continued until the end of the experiment at hour 17. This trend contrasts with the no-drug control (-MF), where septum density plateaued at about 0.00276 septa/µm^2^ approximately 13 hours after inoculation. The trends diverged such that, at 16 hours post inoculation, the septum density of the drug-treated mycelia was 85.0% higher than that of the no-drug control (p<0.0001) (Figure 1, Figure 2).

**Figure 2.**
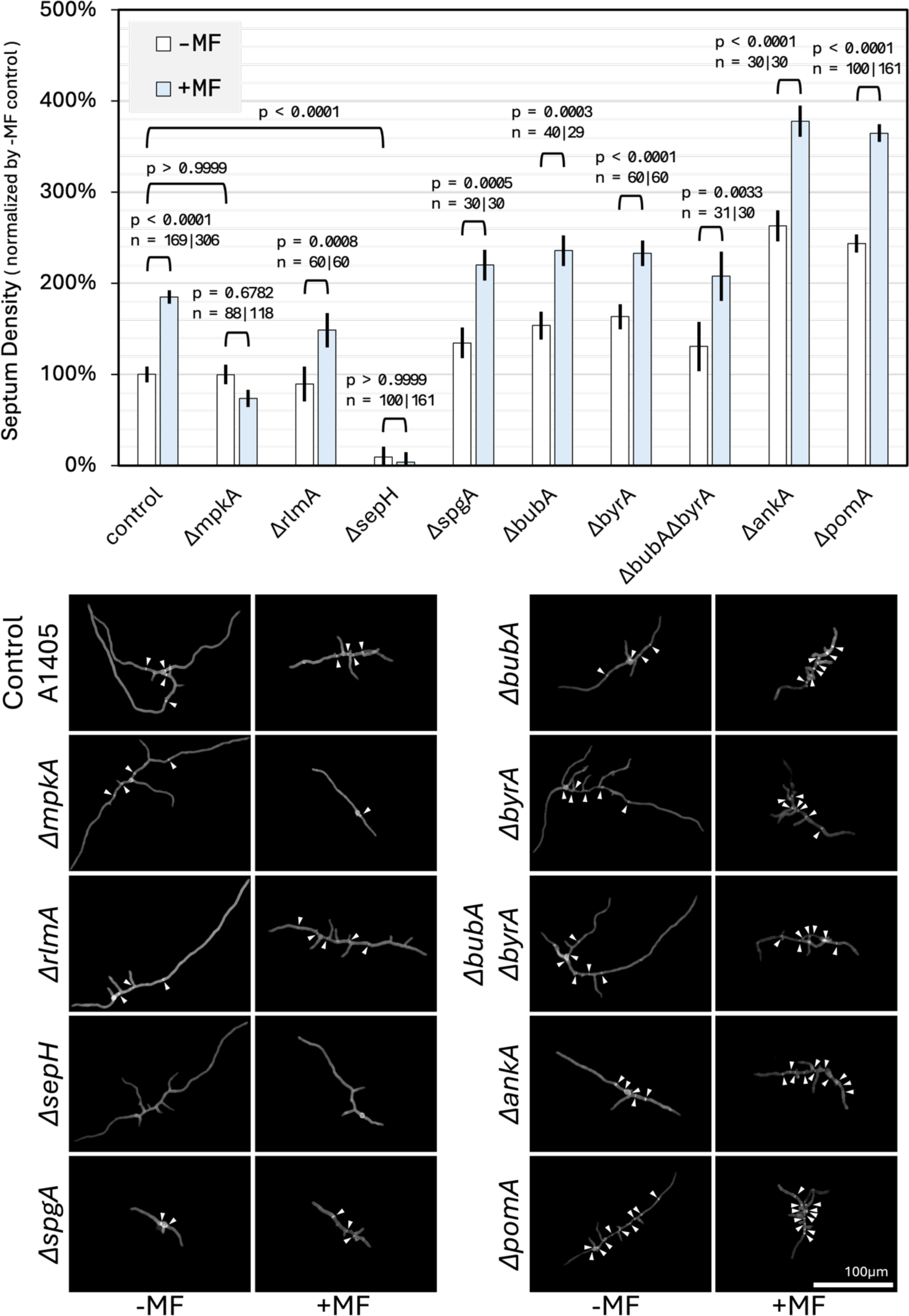
<u>Top</u>: Normalized septum density 16 hours after coverslip inoculation. For shaded bars, cell wall stress was induced with 10 ng/mL micafungin (MF) starting at hour 12. For each −/+ MF pair, the p-value of the contrast is displayed along with the sample size of both bars. Error bars are standard error. <u>Bottom</u>: Selected images represented by bar plot data. White arrowheads highlight septa. Control germlings from Figure 1 are shown again for comparison with knockout germlings.

### SIN is Activated by CWI Via MpkA

Our time-course experiments revealed that 16 hours was the optimal timepoint to run statistical testing between phenotypes (see *Septum-Density Coverslip Assay* in Methods), so in Figure 2, the remaining septum density measurements were made at the 16th hour.

We proceeded with a *ΔmpkA* strain (FGSC A1404)^69,79^ to determine if MpkA, the terminal kinase of CWI, is required for the *wild-type* response to micafungin treatment. In Figure 2, septum density is unresponsive to, and does not increase under, micafungin-induced wall stress (p=0.6782) for the *ΔmpkA* strain. This implies that MpkA is essential to activating the SIN pathway under cell wall stress conditions. The *ΔmpkA* strain also has comparable septum density to the control strain in the no-stress condition, where (p > 0.9999, Figure 2), implying that MpkA is removed from septation regulation in the absence of wall stress. These results highlight that CWI, rather than a parallel upstream pathway, is responsible for SIN activation in response to cell wall stress.

To determine whether the crosstalk occurs downstream of CWI through the transcription factor RlmA, an *ΔrlmA* strain was constructed. Septum density was 66.3% higher under cell wall stress (p=0.0008, Figure 2) than no-stress for the RlmA knockout, which implies the crosstalk signal is not transduced transcriptionally through RlmA regulation and exits CWI through MpkA.

### SIN Receives CWI Signal Through SepH

To identify the entry point of the CWI signal into the SIN pathway, we used a genetic approach and deleted known SIN signaling proteins systematically. We began our investigation with the protein kinase SepH.

In the absence of SepH, septation is significantly reduced, with septum density at 9.3% that of the control strain in the no-stress condition (p<0.0001, Figure 2). This finding is consistent with literature showing that SepH plays a role in the assembly of the septation initiation complex at the spindle pole body^26,28,51,55^. Furthermore, no change in septum density is observed under micafungin-induced cell wall stress conditions in the *ΔsepH* mutant (p>0.9999, Figure 2). These results suggest that SepH is essential for transducing the CWI signal to the SIN pathway and that the signal must enter SIN at or upstream of SepH.

We identified five upstream regulators of SepH in the literature: a small G-Protein, SpgA^26,52^, its two GAPs, BubA and ByrA^30,47^, and two protein kinases, AnkA and PomA^52^. To determine if any of these SIN regulators receive the CWI stress signal, we systematically knocked out the corresponding protein-coding genes of each regulator and measured septum density with, and without, micafungin treatment. Additionally, we created and tested a double knockout of the two GAPs, that regulate SpgA in parallel, to rule out the possibility that the GAPs transduce redundant CWI signals.

In Figure 2, for each of the knockouts *ΔspgA, ΔbubA, ΔbyrA, ΔbubAΔbyrA, ΔankA,* and *ΔpomA,* the *wild-type* response is observed, where septum density is higher under cell wall stress conditions compared to the unstressed condition (all p < 0.01). As no proteins upstream of SepH in the SIN pathway were found to be essential for signal transduction, we propose that SepH serves as the entry point of the CWI stress signal into the SIN pathway.

## DISCUSSION

In this work we discovered that the Septation Initiation Network (SIN) responds to cell wall stress by increasing septum density. Using a genetic approach, we found that the septation response is mediated by a crosstalk signal, where CWI activates SIN during wall stress conditions.

Specifically, we found that the stress signal is transmitted from CWI by MpkA and is received in SIN by SepH (Figure 3). Nominally, septum formation is regulated through the cyclin-dependent kinase (CDK) NimX and is thereby coupled to mitosis and the larger cell cycle^26,80–83^. Our results demonstrate that septum formation can be propagated independently by stress-induced signaling through the CWI kinase MpkA. In this mapping, SIN is shared by both processes, with SepH serving as the junction in Figure 3.

**Figure 3.**
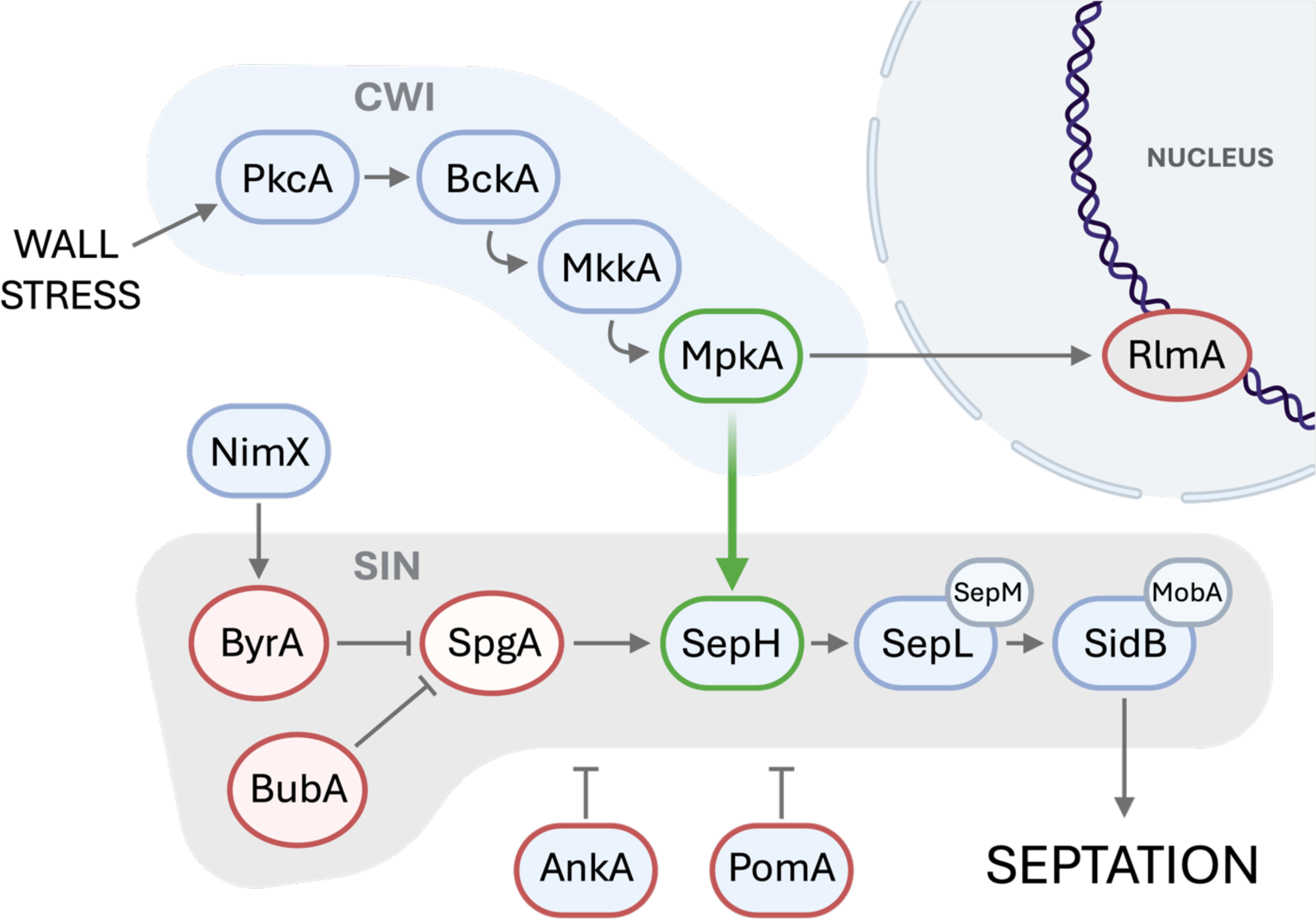
Signal diagram of Cell Wall Integrity (CWI) and Septation Initiation Network (SIN) pathways. Red Border = knockout strain had *wild-type* response to wall stress (increased septum density), Green Border = knockout strain did not change septum density in response to wall stress. Our results suggest that crosstalk is transduced between MpkA (CWI) and SepH (SIN) kinases under cell wall stress conditions, highlighted by the green arrow. Orthologous protein names of other common ascomycete models are listed in Table 1.

We hypothesize that increased septation serves as a survival mechanism for mycelia experiencing weakened cell wall conditions. Mycelia with compromised cell wall integrity are at a higher risk of losing septal compartments due to hyphal injury; increasing the septum density manages this risk by limiting the amount of cytoplasm that can be lost during a cell wall failure event. Our hypothesis is supported by previous findings demonstrating that impaired septation results in significantly lower survival under cell wall stress conditions^23–25^, and furthermore by our findings in Figure S1, and supporting literature^18,23,24^, showing that disabled CWI-SIN crosstalk, through the deletion of *mpkA* or *sepH*, significantly reduces cell wall stress tolerance. Together, these findings provide considerable evidence that the ability to regulate septum density in response to wall stress is critical to survival.

Deletion of the CWI transcription factor RlmA does not result in reduced survival under wall stress (Figure S1), implying that transcription plays a diminished role in surviving echinocandin wall stress.

Our findings in *A. nidulans* have been strikingly consistent with those reported in *A. fumigatus.* Namely, deletion mutants of SIN orthologues *sepH* and *sepL* survive at significantly lower rates when exposed to echinocandin wall stress^23,24^, whereas deletion mutants of SIN orthologues *spgA, bubA,* and *byrA* do not in both *A. fumigatus*^24,25^ and *A. nidulans*^23^ (Figure S1). Septation is also completely eliminated, or considerably reduced, for *ΔsepL* (Figure S2) and *ΔsepH* deletions, respectively, in both species, whereas it is not for knockouts of other SIN regulators^23,24^. These consistencies suggest that the septation response to wall stress is at least conserved broadly across Eurotiales. Furthermore, the conservation of CWI and SIN pathway components across a wide range of fungal species^19,26,48,73,84–88^, raises the possibility that the crosstalk mechanism discovered in this study may, in some semblance, be employed by fungi more broadly to respond to cell wall stress.

To the best of our knowledge, our discovery represents the first time CWI-SIN crosstalk has been described in filamentous ascomycetes and may be applicable to fungi more broadly. Comparative genomic and functional analyses across more diverse fungal species could shed brighter light on the evolutionary origins and functional significance thereof. In particular, the septation response to cell wall stress is likely relevant to clinically significant fungal pathogens. Echinocandins target cell wall synthesis and are widely used to treat mycoses^5,44,89^. Their efficacy is species specific though^5,43^, which our results suggest is due to the CWI-SIN response and how it enables filamentous species to fortify with septa in response to echinocandin stress. Hence, our discovery of CWI-SIN crosstalk advances the molecular understanding of echinocandin pharmacology and will help to inform development of novel fungal treatment strategies to overcome echinocandin resistance in filamentous fungi.

## METHODS

### Strains

A full list of strains used in this study is available in Data Sheet S3.

Our experiments were carried out using strains in the SO451 background (*pyrG89; wA3; argB2; ΔnkuAku70::argB pyroA4; sE15 nirA14 chaA1 fwA1*)^69,79,90^. Several of the kinase knockout strains had already been generated for the Kinase Knockout Library (KKOL) by De Souza et al. (2013)^69^. These strains (*ΔmpkA, ΔsepH, and ΔpomA*) were acquired from FGSC (strain numbers A1404, A1334, and A1371, respectively)^69,79^. We used the KKOL *wild-type* control strain (FGSC A1405) as the control for our experiments that possesses *pyrG* transformed back into SO451^69^. The remaining knockout strains were SO451 transformants generated using a CRISPR-Cas9 and homology directed repair (HDR) workflow.

### Storage and Preparation

Fresh spores were used in all experiments. Fungal strains are stored in phospho-buffered 20% glycerol stocks at −80°C. Fresh spores were produced by growing up banked spores on MAGVS solid media (2% Glucose, 2% Malt Extract, 0.2% Peptone, 1.5% Agar, 0.95M Sucrose, with Vitamins and Hutner’s Trace Elements supplements)^79^. Conidia were harvested from the MAGVS plates using DI water. The spores were then separated from the hyphal mass and conidiophores with vigorous agitation and subsequent filtration through glass wool. Spore concentrations were measured under microscope with a hemocytometer.

### Septum-Density Coverslip Assay

Spores were adhered to coverslips for the duration of the time-course study with concanavalin A (conA) (ThermoFisher J61221). Unless stated otherwise, for all of the steps of the procedure, the same side of the coverslip was always facing in the upward direction to ensure that all of the treatments were performed to the same side of the coverslip. Square coverslips (no. 2, 25 mm) were dusted off with compressed air, then autoclaved. Sterile coverslips were then submerged in a conA solution (100µL 132mM CaCl_2_, 10.7 mg conA in 20 mL PBS) for 30 minutes at 30 °C. They were then dipped in DI water 3 times and placed on Teflon squares inside of a sterile petri dish to dry overnight.

Spores were adhered to the coverslips by pipetting 1 mL of a 5e4 spore/mL solution directly onto the surface of each coverslip. Excess spore solution was pipetted up, leaving the coverslips damp. The spores were left to adhere to the coverslips for 1 hour at 30°C. Then the coverslips were removed from their Teflon squares and submerged in sterile petri dishes filled with 25 mL pre-heated liquid YGV media (0.5% Yeast Extract, 2% Glucose, with Vitamins and Hutner’s Trace Elements supplements)^79^. The petri dishes were then placed in an oven-style incubator set to 28 °C. For the micafungin stress condition, 25 µL of 10 µg/mL micafungin in DI water was added to the petri dish 12 hours after the start of incubation (10 ng/mL final conc). The micafungin was mixed into the media by gently pipetting the media up and down with a 1000 µL pipette.

Enough coverslips were prepared so that one coverslip per timepoint could be harvested for imaging. At each timepoint, a coverslip was removed from the liquid media and submerged for at least 30 minutes in a petri dish containing filtered fixative (3.7% paraformaldehyde, 0.2% Triton X-100, 50mM NaH_2_PO_4_, KOH flakes until clear, to pH 7 with HCl). After fixing, a microscope slide was cleaned with 95% EtOH and 12 µL staining solution (1 mg/mL calcofluor white, 0.5 mg/mL Evan’s blue solution, mixed 1:1 with glycerol, filtered) was pipetted onto the middle of the slide. A coverslip was then removed from the fixative dish, blotting excess fixative onto a sterile Kim-Wipe, and placed face-down on the slide over the drop of stain. Gentle pressure was used to flatten the coverslip and mycelia onto the slide.

Approximately 30 germlings were imaged for each timepoint. Growth area (pixels), number of tips, number of germ tubes, and number of septa were recorded for each germling. Images were taken using an Olympus IX81 microscope equipped with a fluorescent Lumencor SOLA Light Engine and Hamamatsu ORCA-spark camera. Area in pixels was measured using ImageJ’s automatic threshold feature. ImageJ measurements were validated by eye visually and taken with as little manual intervention as possible. Pixels were converted to µm^2^ using a hemocytometer to calibrate the conversion.

In our time-course study, we found that the 16th hour timepoint offered the maximum difference in septum density between the drug and no-drug treatments without a drop-off in in the number of germlings measured per coverslip. At timepoints beyond 16 hours, the germlings became increasingly large and interwoven, leading to a decrease in the number of suitable germlings available to image per timepoint and a bias towards smaller individuals. Having established the 16th hour as the optimal timepoint for analysis, we transitioned to a single-endpoint strategy for the knockout strains, allowing us to compare treatments independent of time. Hence, for each treatment, we measured septum density 16 hours after inoculation. To facilitate comparison in this study, we normalized single-endpoint septum density measurements to the mean no-drug control strain density at the 16th hour timepoint (0.002758 septa/µm^2^).

### Significance Testing

For septum density data in Figure 1 and Figure 2, standard error bars and p-values were calculated with a one-way ANOVA table of a mixed-effects model^91^. To account for variation between experimental batches, the day each datapoint was collected was considered to be a random-effect in the mixed-effects model. The analysis was run in R^92^ using the car^93^, lme4^91^, and emmeans^94^ libraries.

### CRISPR-Cas9 Transformation Procedure

Our transformation protocol was adapted from the protoplasting procedure described in Oakley et al. (2012)^95^ and the CRISPR-Cas9 mediated transformation procedure described in van Rhijn et al. (2020)^96^.

All plastic- and glass-ware was pre-rinsed with DI water (nuclease free when required) to remove any lipids and detergents that may lyse protoplasts, then 1e8 SO451 spores were incubated in 20 mL non-selective, YGV+UU, medium (0.5% Yeast Extract, 2% Glucose, with Vitamins and Hutner’s Trace Elements, Uridine, and Uracil supplements)^79^ in a 50 mL flask and incubated for 16 hours at 30 °C in a shaker-incubator set to 120 rpm. Mycelia were harvested by filtering the entire flask contents through sterile Miracloth in a Buchner funnel. The mycelia were washed in 2-3 volumes DI water and then raked into a central mass using a flamed spatula. The mycelia were then transferred into a clean 50 mL flask containing 16 mL protoplasting medium.

Protoplasting media was prepared freshly an hour before harvesting mycelia. The 2X protoplasting solution contains 2 g VinoTaste Pro (Novozymes) in 10 mL KCl/Citrate Solution (1.1 M KCl, 0.1 M Citric Acid, pH 5.8 with KOH). Lipids were removed by centrifugation at 4 °C, 5000 x g for 15 minutes followed by extraction of the lipid-free bottom layer by careful pipetting. The solution was filter-sterilized through a 0.22 µm syringe filter (MillexGV) that was pre-washed with a volume of KCl/Citrate solution. The 2X protoplasting solution was then combined 1:1 with 1X non-selective YGV+UU medium to make protoplasting media.

Mycelia were incubated for 1.5 hours in protoplasting media at 30 °C and 100 rpm. During incubation, mycelia were broken up at 10-minute intervals by pipetting the solution up and down with a sterile transfer pipette.

After incubation, the entire flask was chilled on ice for 5 minutes, then the entire contents of the flask were carefully pipetted down on top of a cushion of 1.2 M sucrose (appx 15 mL) pre-chilled to 4 °C in a 50 mL centrifuge tube. Care was taken to keep the two liquid layers separated in the centrifuge tube, which was then spun for 10 minutes at 2800 x g, 4 °C. After spinning, the protoplasts were resting on top of the sucrose layer and were extracted carefully using a sero-pipette. Isolated protoplasts were washed 3 times by repeated cycles of centrifugation at 2400 x g for 3 min at room temperature followed by resuspension in 1 mL 0.6 M KCl. A fourth wash cycle was carried out using a 0.6 M KCl, 50 mM CaCl_2_ solution. The protoplast concentration of the washed solution was measured by hemocytometer and was either diluted or concentrated by centrifugation to 5e6 protoplasts per mL.

The CRISPR/Cas9 ribonucleoprotein (RNP) complex was prepared immediately before transformation by adding in order: 22 µL HEPES Buffer (20 mM HEPES Free Acid, 150 mM KCl, pH to 7.5 with KOH), 0.62 µL Cas9 Dilution Buffer (supplied with Cas9 Plus enzyme), 0.88 µL 1.7 µg/µL Cas9 Plus (Sigma-Aldrich CAS9PL), 2 µL Duplex Buffer (IDT 11-01-03-01), 0.5 µL 100 µM 5p sgRNA, 0.5 µL 100 µL 3p sgRNA. The solution was vortexed, spun, and allowed to incubate at room temperature for 5 minutes.

The transformation solution was assembled by adding, in order, to the assembled RNP complex: 200 µL 5e6 / mL protoplast solution, 2-10 µg linear DNA repair cassette (max vol 50 µL), 25 µL 60% PEG solution (60% w/v PEG 3350, 50 mM CaCl_2_, 450 mM Tris-HCl, pH 7.5 with KOH). The solution was gently vortexed and incubated on ice for 50 minutes. At 10-15 minute intervals the solution was gently vortexed to maintain homogeneity. CRISPR sgRNAs and DNA oligos used to produce the deletion cassettes can be found in Data Sheet S4.

To prepare the 60% PEG solution, 6 g PEG 3350 (Sigma-Aldrich 202444) was carefully melted in a small glass bottle with constant shaking while moving in and out of the heat of a flame. Care was taken not to overheat the PEG. Melted PEG is a clear solution; any browning is indication that the PEG was burned and that it should be discarded. Using 3 mL DI water, 0.074 g CaCl_2_ was added to the melted PEG. Finally, 1.34 mL 2.7 M Tris-HCl and 0.327 mL 2.7 M Tris-Base were added to the solution and mixed over the flame for a final pH of approximately 7.5. The solution was autoclaved, mixed one more time, and then run through a pre-rinsed Millex GP syringe filter.

Membrane permeation was achieved through heat shock. After 50 minutes of incubation on ice, 1.2 mL warm (∼45 °C) 60% PEG solution was added to the transformation solution and rapidly mixed by pipetting up and down and then by gentle vortexing. The solution was allowed to incubate for 20 minutes at room temperature, vortexing at 10 minute intervals to maintain homogeneity.

Plates with MM+VS selective media plates (1% Glucose, 1.5% Agar, 0.05% MgSO_4_, 0.95M Sucrose, 0.6% NaNO3, 0.152% KH_2_PO_4_, 0.052% KCl, with Vitamins and Hutner’s Trace Elements supplements) were each inoculated with 50 µL transformation solution and incubated at 28°C until conidiation was observed. Colonies were then picked and re-plated for genotyping by diagnostic PCR (Figure S6)

## DATA AVAILABILITY

Data presented in this work are available in the figures and supplemental files. Strains used in this study have been retained and are available from FGSC^79^ and/or upon request to the corresponding author.

## SUPPLEMENTAL MATERIALS

**Figure S1-.**
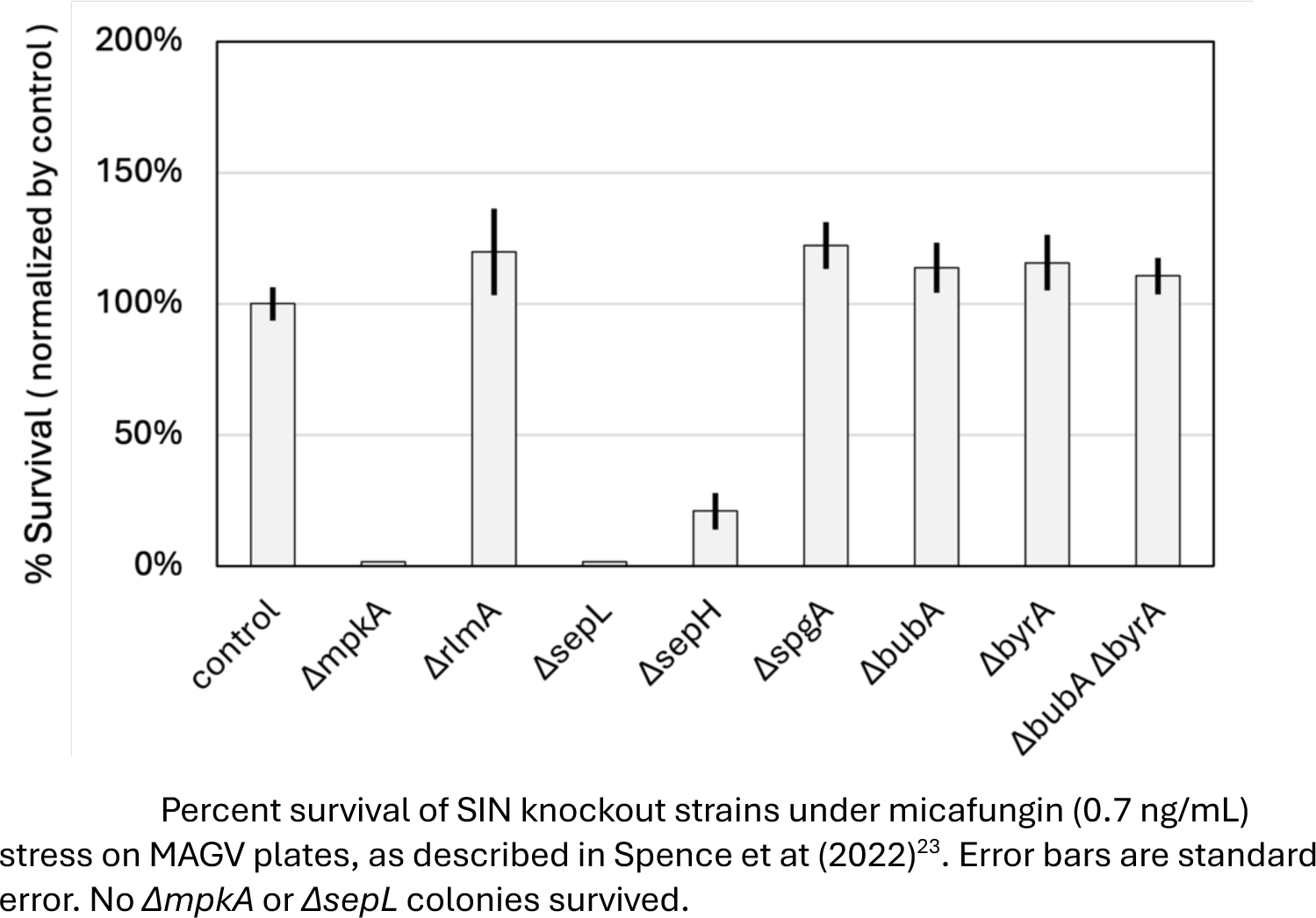
Echinocandin Survival

**Figure S2-.**
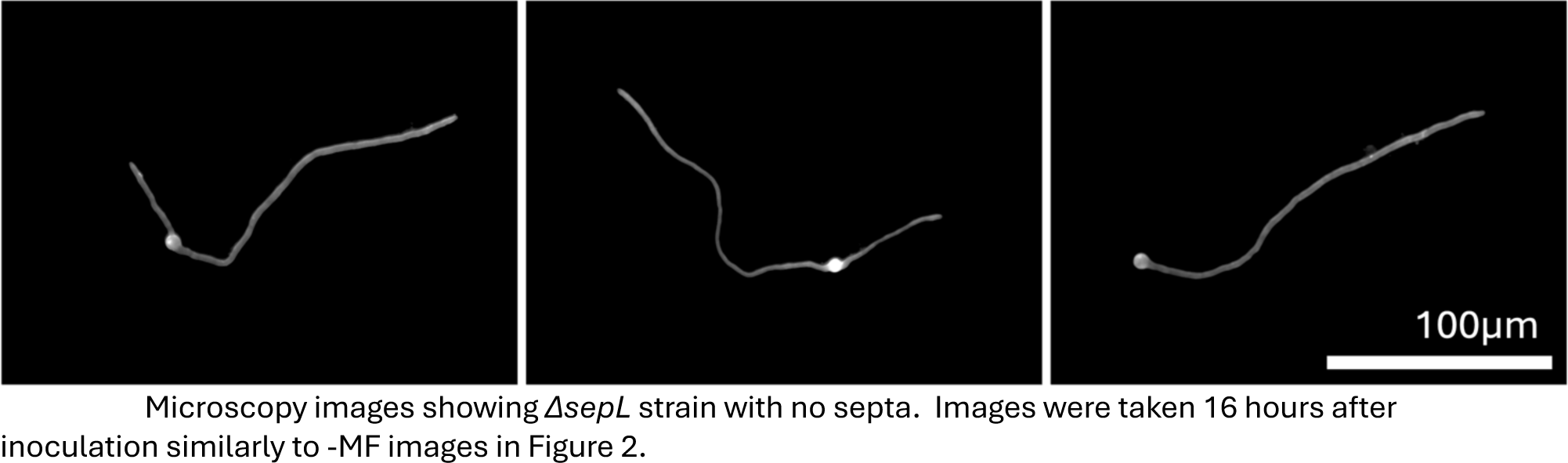
SepL Deletion Images

Data Sheet S3 - Strain List

| Referred to as | Source | Parent | Full Genotype |
| --- | --- | --- | --- |
| SO451/Parent | FGSC, DeSouza et al. 2013 |  | *pyrG89; wA3; argB2; ΔnkuAku70::argB pyroA4; sE15 nirA14 chaA1 fwA1* |
| Control | FGSC, DeSouza et al. 2013 | SO451 | *pyrG<sup>Af</sup>; pyrG89; wA3; argB2; ΔnkuAku70::argB pyroA4; sE15 nirA14 chaA1 fwA1* |
| *ΔsepH* | FGSC, DeSouza et al. 2013 | SO451 | *ΔsepH::pyrG<sup>Af</sup>; pyrG89; wA3; argB2; ΔnkuAku70::argB pyroA4; sE15 nirA14 chaA1 fwA1* |
| *ΔpomA* | FGSC, DeSouza et al. 2013 | SO451 | *ΔpomA::pyrG<sup>Af</sup>; pyrG89; wA3; argB2; ΔnkuAku70::argB pyroA4; sE15 nirA14 chaA1 fwA1* |
| *ΔmpkA* | FGSC, DeSouza et al. 2013 | SO451 | *ΔmpkA::pyrG<sup>Af</sup>; pyrG89; wA3; argB2; ΔnkuAku70::argB pyroA4; sE15 nirA14 chaA1 fwA1* |
| *ΔspgA* | This Study | SO451 | *ΔspgA::pyrG<sup>Af</sup>; pyrG89; wA3; argB2; ΔnkuAku70::argB pyroA4; sE15 nirA14 chaA1 fwA1* |
| *ΔbubA* | This Study | SO451 | *ΔbubA::pyrG<sup>Af</sup>; pyrG89; wA3; argB2; ΔnkuAku70::argB pyroA4; sE15 nirA14 chaA1 fwA1* |
| *ΔbyrA* | This Study | SO451 | *ΔbyrA::pyrG<sup>Af</sup>; pyrG89; wA3; argB2; ΔnkuAku70::argB pyroA4; sE15 nirA14 chaA1 fwA1* |
| *ΔankA* | This Study | SO451 | *ΔankA::pyrG<sup>Af</sup>; pyrG89; wA3; argB2; ΔnkuAku70::argB pyroA4; sE15 nirA14 chaA1 fwA1* |
| *ΔbubAΔbyrA* | This Study | AGD002 | *ΔbyrA::pyroA<sup>Af</sup>; ΔbubA::pyrG<sup>Af</sup>; pyrG89; wA3; argB2; ΔnkuAku70::argB pyroA4; sE15 nirA14 chaA1 fwA1* |
| *ΔrlmA* | This Study | SO451 | *ΔrlmA::pyrG<sup>Af</sup>; pyrG89; wA3; argB2; ΔnkuAku70::argB pyroA4; sE15 nirA14 chaA1 fwA1* |

Data Sheet S4 - Deletion Cassette Oligos

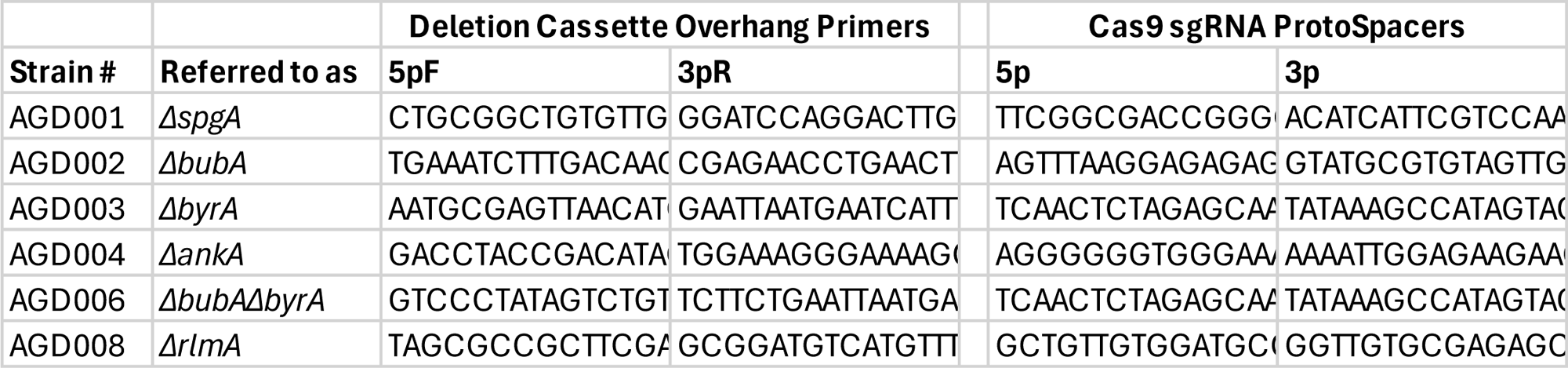

Data Sheet S5 - Diagnostic PCR Oligos

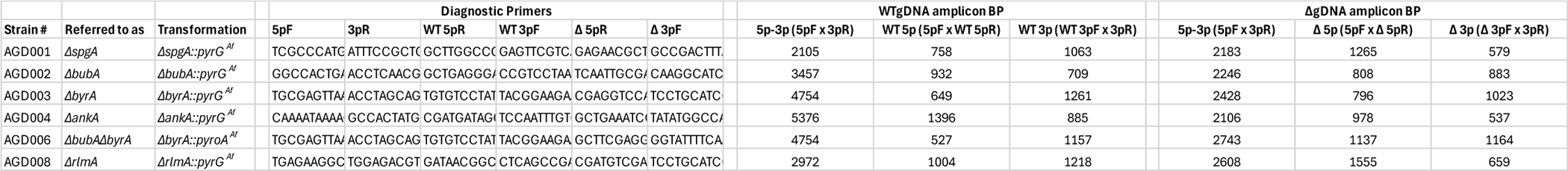

| Strain # | Referred to as | Transformation | Diagnostic Primers |  |  |  |  |  | WTgDNA amplicon BP |  |  | ΔgDNA amplicon BP |  |  |
| --- | --- | --- | --- | --- | --- | --- | --- | --- | --- | --- | --- | --- | --- | --- |
|  |  |  | 5pF | 3pR | WT 5pR | WT 3pF | Δ 5pR | Δ 3pF | 5p-3p (5pF x 3pR) | WT 5p (5pF x WT 5pR) | WT 3p (WT 3pF x 3pR) | 5p-3p (5pF x 3pR) | Δ 5p (5pF x Δ 5pR) | Δ 3p (Δ 3pF x 3pR) |
| AGD001 | *ΔspgA* | *ΔspgA::pyrG<sup>Af</sup>* | TCGCCCATC | ATTTCGCCT | GCTTGCCC | GAGTTCGTC | GAGAACGCT | GCCGACTTT | 2105 | 758 | 1063 | 2183 | 1265 | 579 |
| AGD002 | *ΔbubA* | *ΔbubA::pyrG<sup>Af</sup>* | GGCCACTG | ACCTCAACG | GCTGAGGG | CCGTCCTAA | TCAATTGCG | CAAGGCATC | 3457 | 932 | 709 | 2246 | 808 | 883 |
| AGD003 | *ΔbyrA* | *ΔbyrA::pyrG<sup>Af</sup>* | TGCGAGTTA | ACCTAGCAG | TGTGTCCTA | TACGGAAGA | CGAGGTCC | TCCTGCATC | 4754 | 649 | 1261 | 2428 | 796 | 1023 |
| AGD004 | *ΔankA* | *ΔankA::pyrG<sup>Af</sup>* | CAAAATAAAA | GCCACTATG | CGATGATAG | TCCAATTGT | GCTGAAATC | TATATGGCC | 5376 | 1396 | 885 | 2106 | 978 | 537 |
| AGD006 | *ΔbubAΔbyrA* | *ΔbyrA::pyroA<sup>Af</sup>* | TGCGAGTTA | ACCTAGCAG | TGTGTCCTA | TACGGAAGA | GCTTCGAGC | GGTATTTCA | 4754 | 527 | 1157 | 2743 | 1137 | 1164 |
| AGD008 | *ΔrlmA* | *ΔrlmA::pyrG<sup>Af</sup>* | TGAGAAGGC | TGGAGACGT | GATAACGGC | CTCAGCCG | CGATGTCGA | TCCTGCATC | 2972 | 1004 | 1218 | 2608 | 1555 | 659 |

**Figure S6-.**
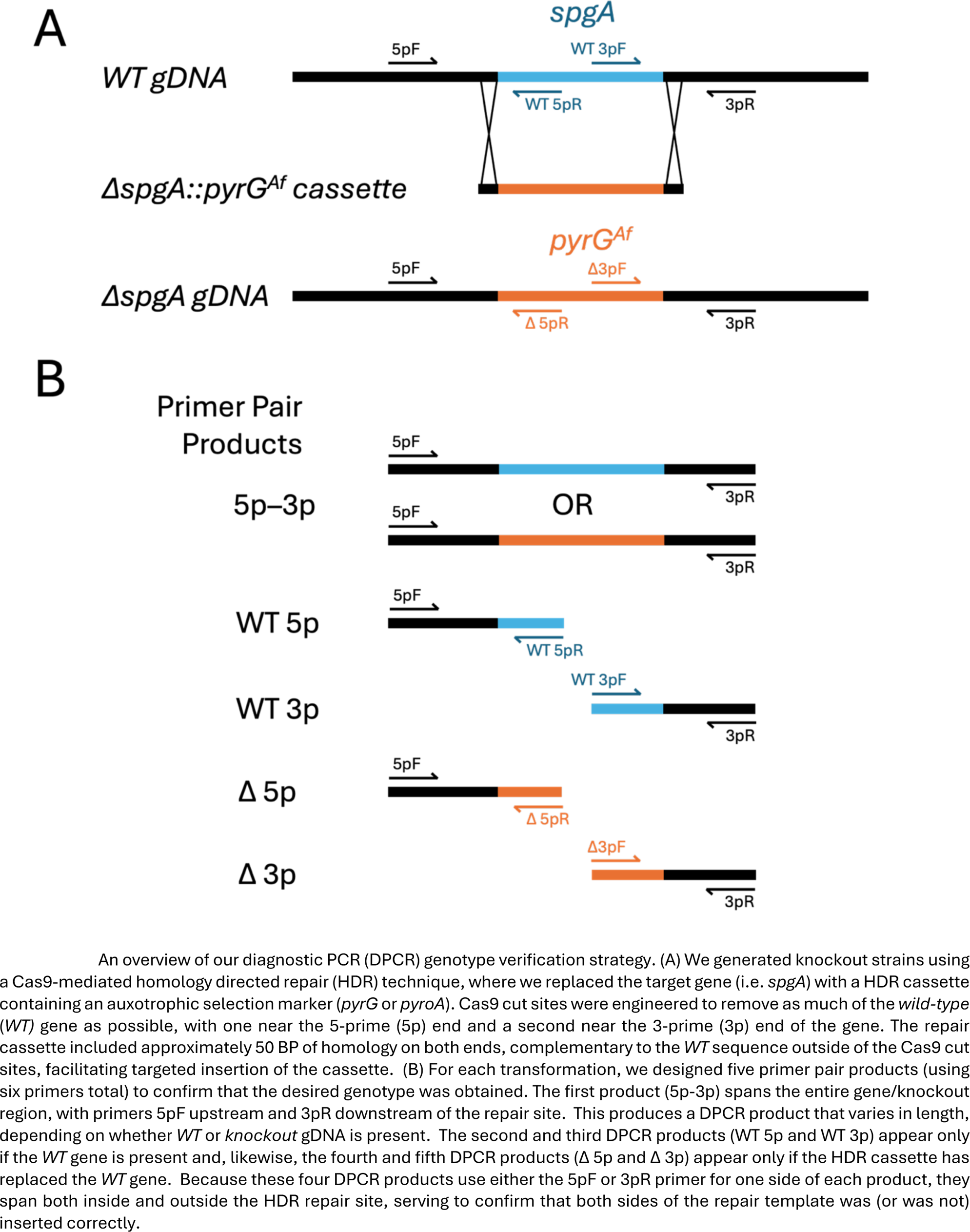
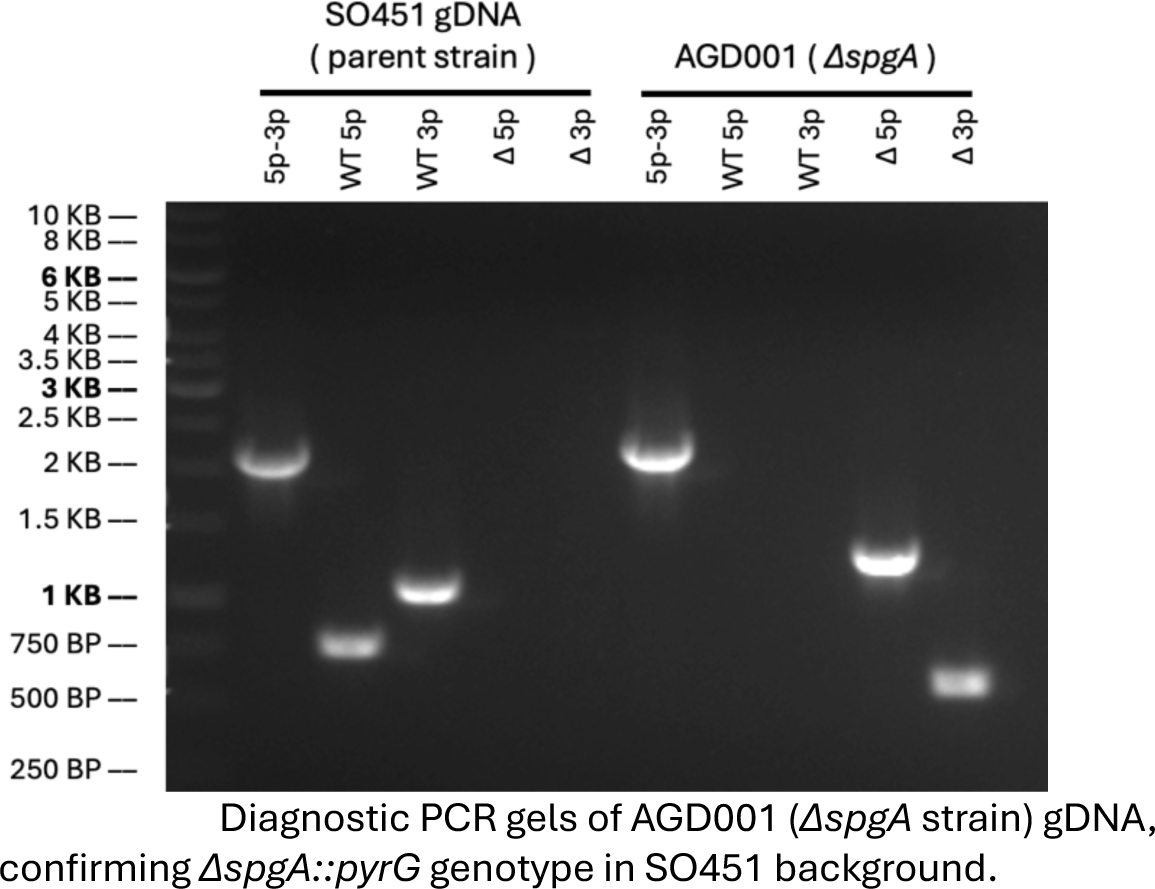
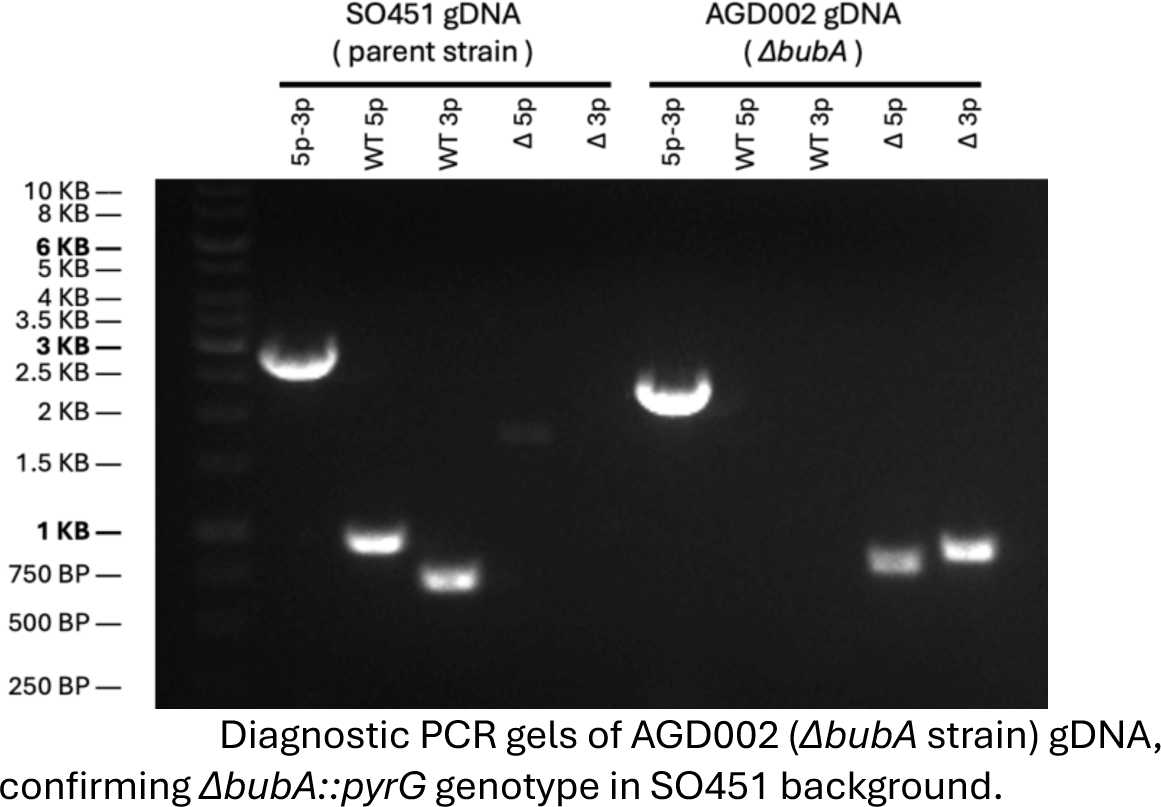
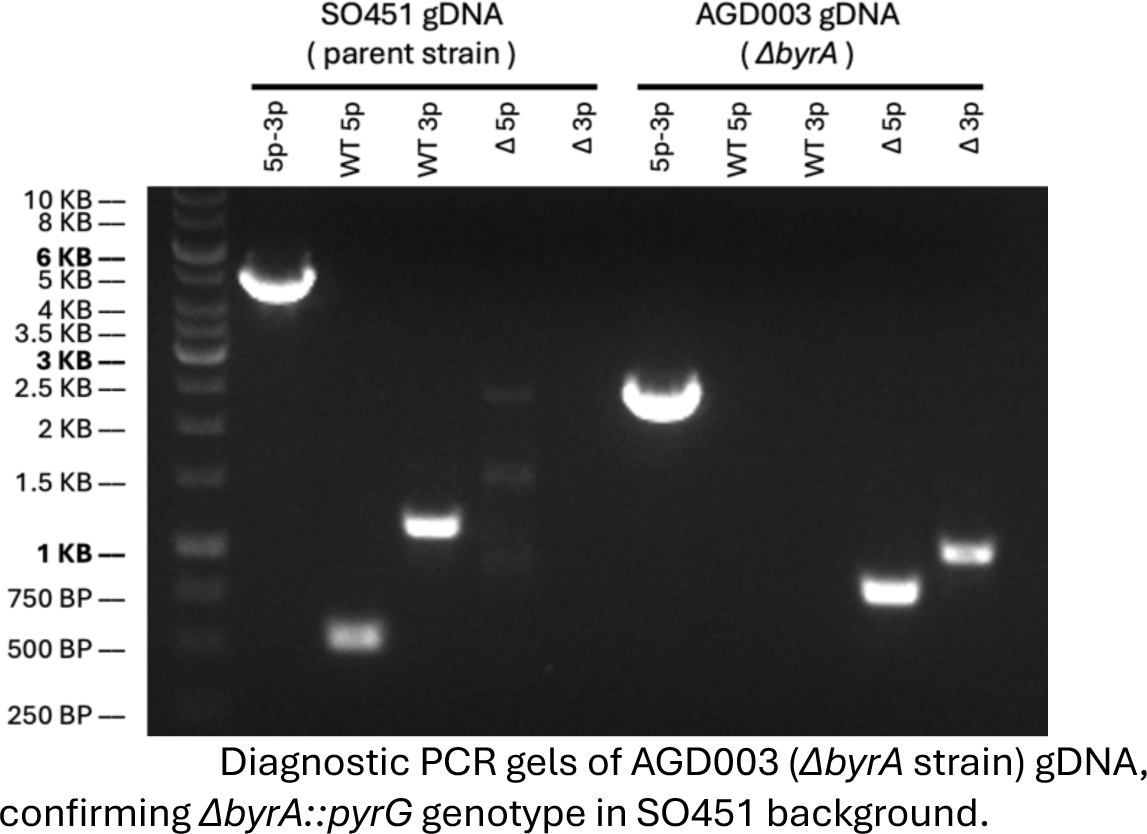
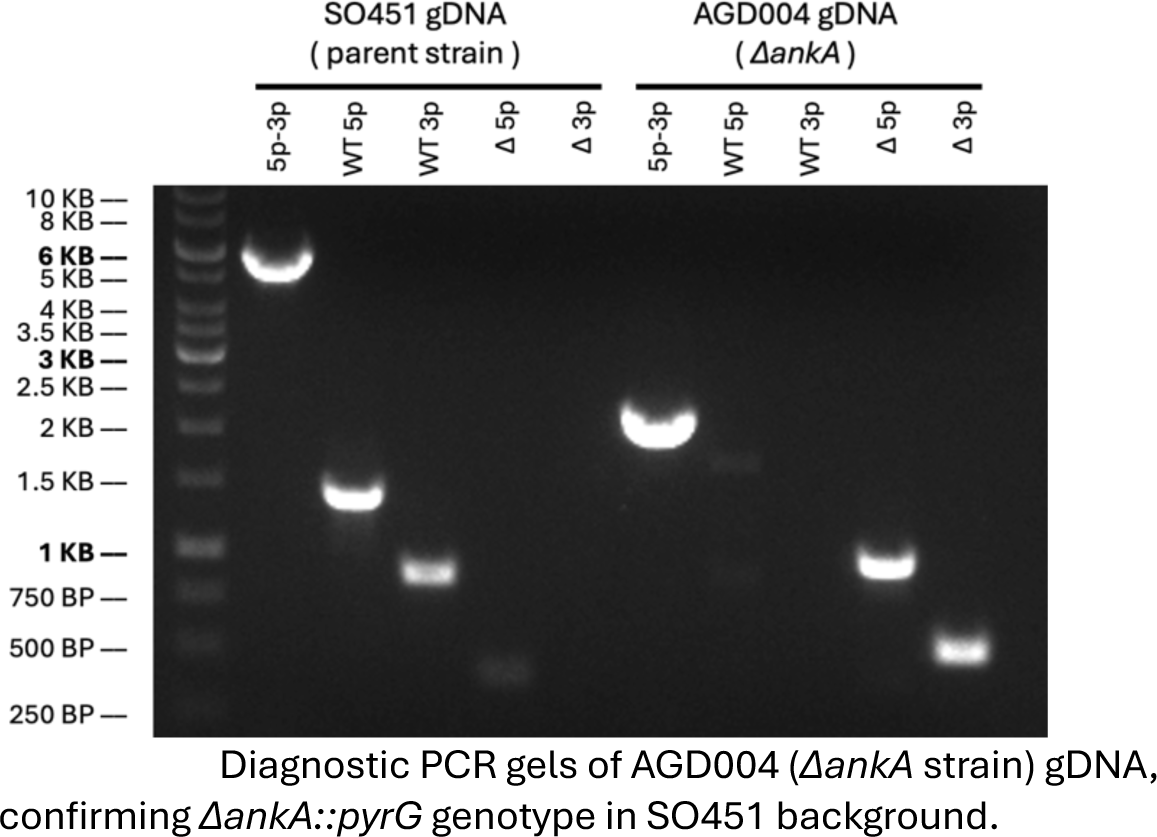
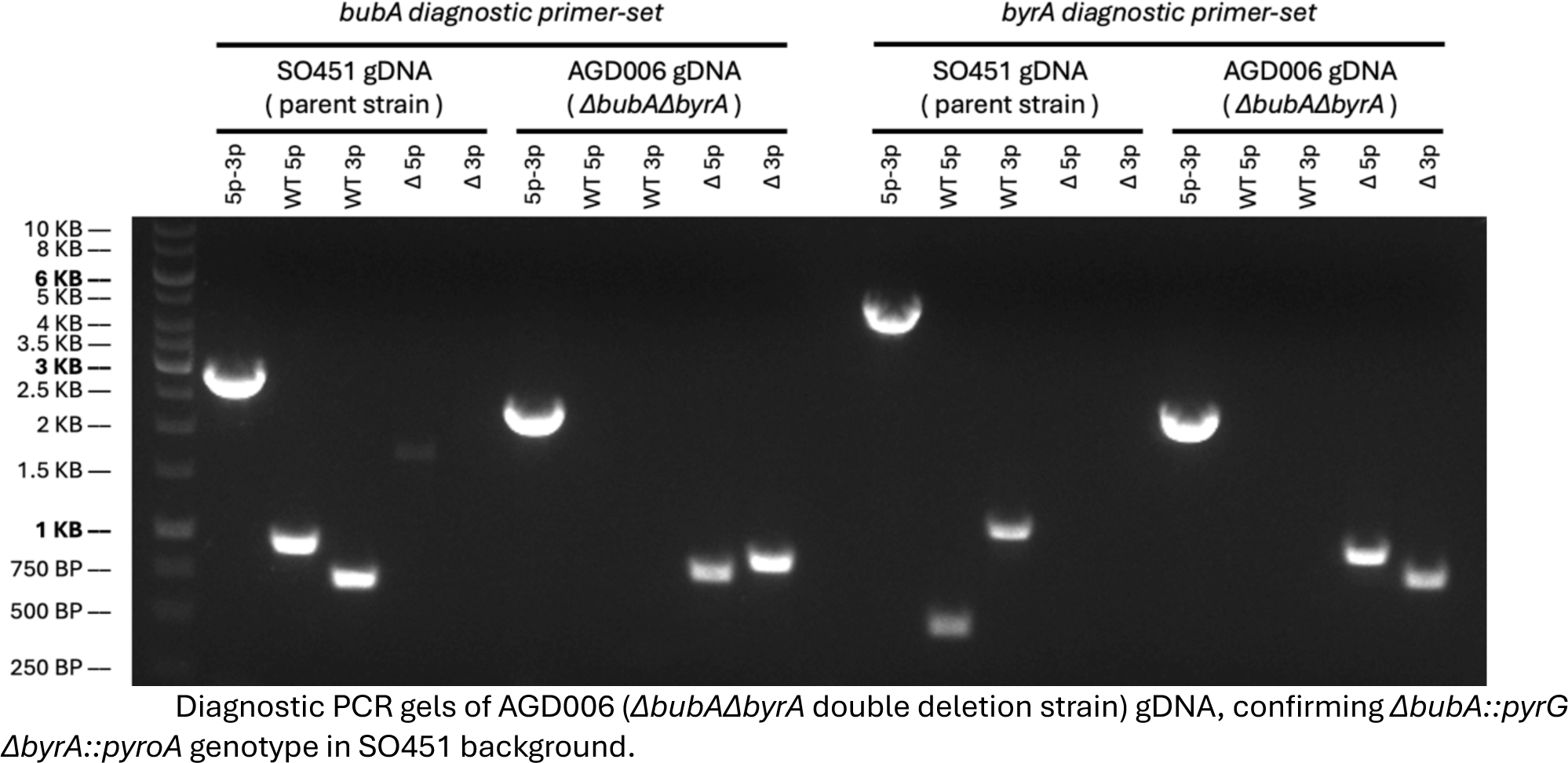
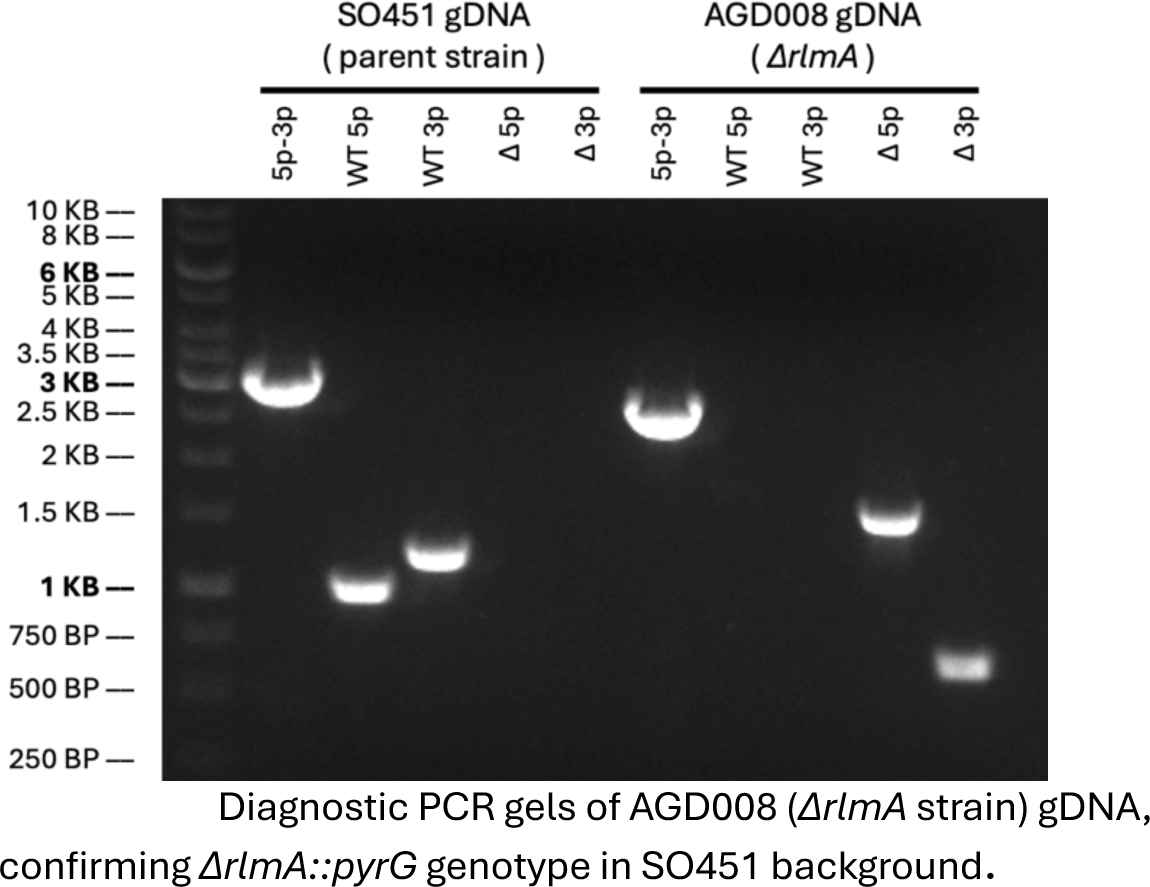
Diagnostic PCR Gels

